# 30S subunit recognition and G1405 modification by the aminoglycoside-resistance 16S ribosomal RNA methyltransferase RmtC

**DOI:** 10.1101/2023.03.13.532395

**Authors:** Pooja Srinivas, Meisam Nosrati, Natalia Zelinskaya, Debayan Dey, Lindsay R. Comstock, Christine M. Dunham, Graeme L. Conn

## Abstract

Acquired ribosomal RNA (rRNA) methylation has emerged as a significant mechanism of aminoglycoside resistance in pathogenic bacterial infections. Modification of a single nucleotide in the ribosome decoding center by the aminoglycoside-resistance 16S rRNA (m^7^G1405) methyltransferases effectively blocks the action of all 4,6-deoxystreptamine ring-containing aminoglycosides, including the latest generation of drugs. To define the molecular basis of 30S subunit recognition and G1405 modification by these enzymes, we used a *S*-adenosyl-L-methionine (SAM) analog to trap the complex in a post-catalytic state to enable determination of an overall 3.0 Å cryo-electron microscopy structure of the m^7^G1405 methyltransferase RmtC bound to the mature *Escherichia coli* 30S ribosomal subunit. This structure, together with functional analyses of RmtC variants, identifies the RmtC N-terminal domain as critical for recognition and docking of the enzyme on a conserved 16S rRNA tertiary surface adjacent to G1405 in 16S rRNA helix 44 (h44). To access the G1405 N7 position for modification, a collection of residues across one surface of RmtC, including a loop that undergoes a disorder to order transition upon 30S subunit binding, induces significant distortion of h44. This distortion flips G1405 into the enzyme active site where it is positioned for modification by two almost universally conserved RmtC residues. These studies expand our understanding of ribosome recognition by rRNA modification enzymes and present a more complete structural basis for future development of strategies to inhibit m^7^G1405 modification to re-sensitize bacterial pathogens to aminoglycosides.

**Significance:** Increasing prevalence of bacterial antibiotic resistance threatens our ability to treat bacterial infections and with it, many other facets of modern healthcare. For the ribosome-targeting aminoglycoside antibiotics, diverse pathogenic bacteria have acquired ribosomal RNA (rRNA) methyltransferase enzymes that confer exceptionally high-level resistance through site-specific modification of the drug binding site. Here, we define the molecular basis for ribosomal substrate recognition and modification by an enzyme (RmtC) representing the most clinically prevalent methyltransferase family. Specifically, RmtC exploits a conserved rRNA surface for binding and induces significant disruption of the rRNA structure to capture the target nucleotide for modification via a “base flipping” mechanism. These insights also present a platform for methyltransferase inhibitor development to extend usefulness of aminoglycoside antibiotics.

## Introduction

With rising resistance among pathogenic bacteria against all current antibiotic classes, efforts to define and counter such resistance mechanisms are of critical importance (1, 2). Aminoglycosides are an essential class of ribosome-targeting antibiotics, with activity against both Gram-positive and Gram-negative bacteria (3). However, the efficacy of these drugs is challenged by multiple resistance mechanisms including drug efflux, drug modification by aminoglycoside modifying enzymes, and ribosomal target site alteration via mutation or chemical modification. While drug modification is currently the most widespread cause of clinical aminoglycoside resistance (2), an increasingly prevalent resistance mechanism in multiple human pathogens is the expression of 16S ribosomal RNA (rRNA) methyltransferase enzymes capable of modifying the nucleobase of one of two 16S rRNA nucleotides (G1405 or A1408, in *Escherichia coli* numbering) in the drug binding site on the mature 30S subunit (4-6). With their global dissemination, such enzymes have the potential to confer widespread and high-level resistance to essentially all clinically important aminoglycosides.

Aminoglycoside antibiotics typically act by reducing the fidelity of decoding or inhibiting tRNA movement through the ribosome during translation (7-10). Two 16S rRNA nucleotides, A1492 and A1493, flip out of helix 44 (h44) to sample the mRNA codon-transfer RNA (tRNA) anticodon interaction (11, 12), in a conformation that is normally only stably adopted for a cognate mRNA-tRNA pairing (13). However, aminoglycoside binding to h44 immediately adjacent to A1492 and A1493 promotes adoption of the flipped-out conformation of these bases, allowing a non-cognate tRNA anticodon to be misread as cognate and thus incorporation of the incorrect amino acid during protein synthesis. Methylation of G1405 or A1408 within the h44 aminoglycoside binding site to produce m^7^G1405 or m^1^A1408, respectively (14), blocks drug binding and thus the resulting effect on translation.

The aminoglycoside-resistance 16S rRNA methyltransferases were originally identified in drug biosynthesis gene clusters of aminoglycoside-producing actinomycetes where they prevent self-intoxication (3). Pathogenic bacteria have acquired these genes, but sequence identity across species is moderate (∽25-30%). The encoded Class I *S*-adenosyl-L-methionine (SAM)–dependent methyltransferases are functionally divided into two subfamilies, each specific to either the m^7^G1405 or m^1^A1408 modification. Currently, the m^7^G1405 rRNA methyltransferases, which include ArmA and RmtA-H, are globally disseminated and represent the greater clinical challenge with prevalence rates between 3% and 27% among aminoglycoside-resistant Gram-negative infections in hospitals world-wide (2, 6, 14-16). These rates are likely to continue to rise as the 16S rRNA methyltransferase genes are readily transferable between bacterial species via horizontal gene transfer mechanisms (6).

The structures of several m^7^G1405 and m^1^A1408 rRNA methyltransferases have been determined, revealing the high structural conservation within each subfamily, and elucidating the basis of SAM cosubstrate recognition (17-23). However, despite their adjacent target sites and likely significant overlapping 30S binding surface (24), the m^7^G1405 and m^1^A1408 rRNA methyltransferase subfamilies differ extensively in their appendages to the Class I methyltransferase core fold that control substrate recognition. The Arm/Rmt (m^7^G1405) family have an 80-105 residue N-terminal domain composed of two α-helical subdomains (N1 and N2) (21, 22, 24), whereas the m^1^A1408 methyltransferases, such as NpmA, have a short N-terminal β-hairpin and longer internal extensions between β-strands β5 and β6 (β5/β6-linker), and β6 and β7 (β6/β7-linker) (17, 18). Structural and functional studies of NpmA have defined the process of substrate recognition and m^1^A1408 modification (23, 25), but there is currently no high-resolution structure of any m^7^G1405 rRNA methyltransferase bound to the 30S subunit, significantly limiting our understanding of this clinically more important enzyme subfamily.

Here, we report a cryo-electron microscopy structure of the m^7^G1405 rRNA methyltransferase RmtC bound to the *E. coli* 30S ribosomal subunit enabled by use of a SAM analog that traps the complex in a post-catalytic state. This structure allows rationalization of prior studies from our laboratory and others on RmtC and related enzymes (21, 22, 24), as well as guiding new functional analyses to define the critical interactions that facilitate 30S subunit recognition and induce base flipping of G1405 for modification via a significant distortion of h44.

## Results

### Structure of RmtC bound to the mature *E. coli* 30S ribosomal subunit

*E. coli* 30S subunits and recombinant RmtC were purified as previously described (24). A SAM analog, N-mustard 6 (NM6), that is transferred in its entirety to N^7^ of G1405 by the enzymatic action of RmtC, was used to trap the 30S-RmtC complex in an immediately post-catalytic state (**Supplemental Figure S1**). This complex was then used to determine an overall 3.0-Å cryo-EM structure of RmtC bound to the 30S subunit (**Figure 1, Supplemental Figures S2** and **S3**, and **Table 1**). The resolution of RmtC varies between 3.0-8.0 Å, consistent with its location on the surface of the 30S subunit; however, the surface of RmtC that interacts with the 30S subunit shows the highest resolution (∽3.0-4.0 Å; **Supplemental Figure S3F**,**G**), allowing detailed insight into enzyme-substrate recognition. In this structure, RmtC is positioned with its active site centered over h44 with additional contacts made to helices h24, h27 and h45 which form the adjacent highly conserved 16S rRNA surface of the mature 30S subunit (**Figures 1C-F**).

**Figure 1.**
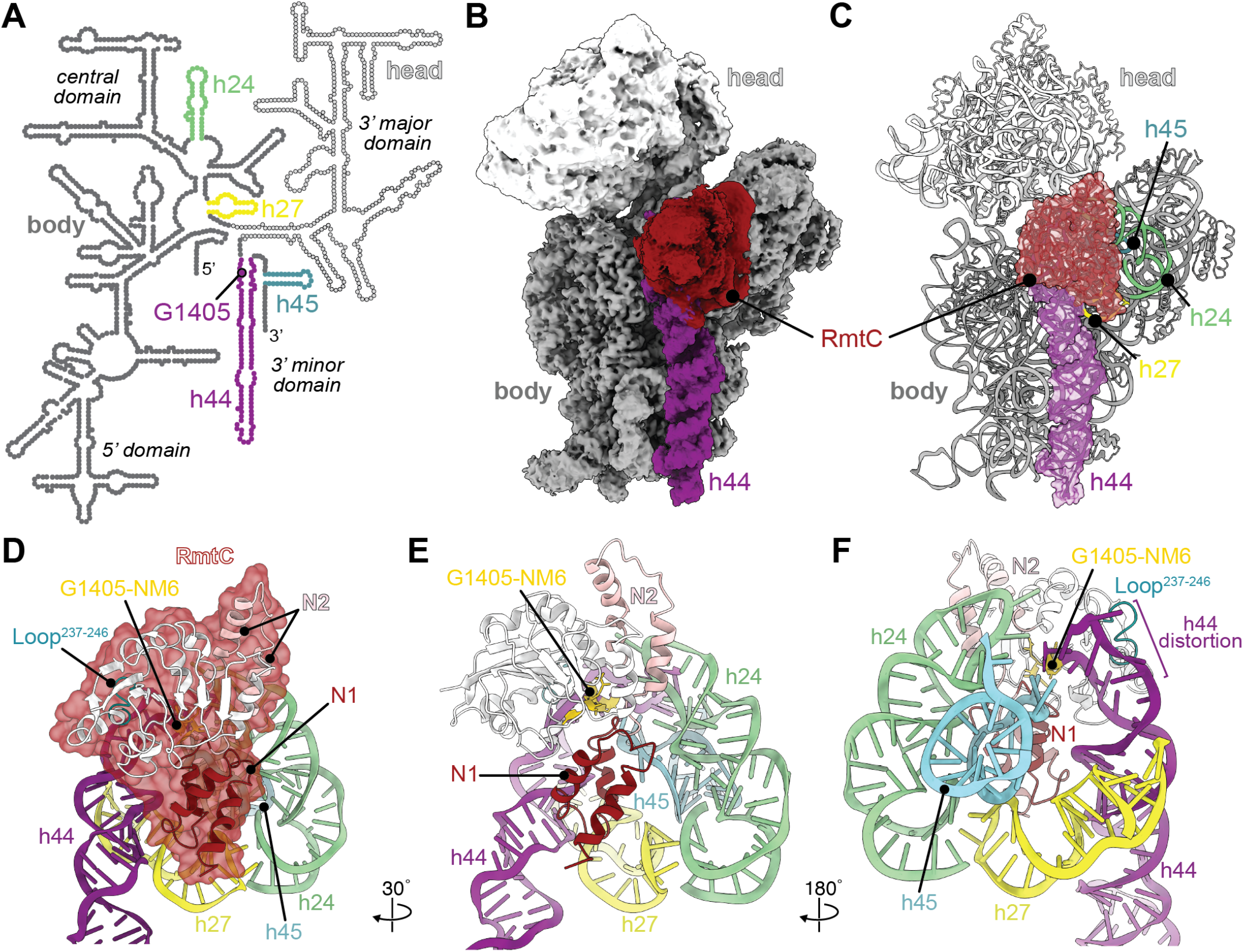
RmtC engages with a conserved 16S rRNA tertiary surface. **A**. Secondary structure of the *E. coli* 16S rRNA, highlighting the modified nucleotide G1405 within h44 (purple), and other rRNA helices, h24 (green), h27 (yellow), and h45 (cyan), that comprise the conserved 16S rRNA surface recognized by RmtC. **B**. Postprocessed cryo-EM map of RmtC (red) bound to the 30S subunit with h44 (purple), body (dark gray) and head (white) domains indicated. **C**. Model of the 30S-RmtC complex, with RmtC and h44 shown as cartoon with a semi-transparent surface representation. **D**. RmtC (red semi-transparent surface) docks on the 30S subunit via interactions with helices h24, h27, h44, and h45 contacts made by both N-terminal subdomains (N1, dark red and N2, pink), and the C-terminal domain (white), including the Loop^237-246^ region (teal). **E** and **F**, Two additional views of RmtC on the 16S rRNA surface.

**Table 1.** Cryo-EM data collection, 30S-RmtC Complex (PDB 8GHU and EMDB 40051) refinement, and validation statistics.

|  | Post-Processed Map | Sharpened Map |
| --- | --- | --- |
| *Data collection* |  |  |
| Magnification | 29,000 | — |
| Voltage (kV) | 300 | — |
| Electron exposure (e<sup>-</sup>/Å<sup>2</sup>) | 51 | — |
| Defocus range (μm) | -0.5 to -3.5 | — |
| Pixel size (Å) | 0.7983 | — |
| Symmetry imposed | C1 | — |
| Initial particle images (no.) | 139,340 | — |
| Final particle images (no.) | 129,736 | — |
| Map resolution (FSC<sub>0.143</sub>) (Å) | 3 | 3.3 |
| Map resolution range (Å) | 2.8-8.0 | — |
| *Refinement* |  |  |
| Model resolution (FSC<sub>0.143</sub>) (Å) | 3 | 3.4 |
| CC<sub>mask</sub> | 0.70 | 0.75 |
| Model resolution range (Å) | 2.6 – 6.6 | 3.3 – 5.2 |
| Map sharpening *B* factor (Å<sup>2</sup>) | -69.8 | -135 |
| Model composition |  |  |
| Nonhydrogen atoms | 52,672 | — |
| Protein Residues | 2490 | — |
| Nucleotides | 1538 | — |
| Ligands | 1 | — |
| Mg<sup>2+</sup> | 83 | — |
| ADP (B-factors) |  |  |
| Protein (min/max/mean) | 1.17/171.26/10.26 | — |
| Nucleotide (min/max/mean) | 0.00/20.00/5.51 | — |
| Ligand (min/max/mean) | 11.89/11.89/11.89 | — |
| R.M.S. deviations |  |  |
| Bond lengths (Å) | 0.004 | — |
| Bond angles (°) | 0.654 | — |
| *Validation* |  |  |
| MolProbity score | 2.01 | — |
| Clashscore | 8.09 | — |
| Rotamer Outliers (%) | 0.48 | — |
| Ramachandran plot |  |  |
| Favored (%) | 89.23 | — |
| Allowed (%) | 10.69 | — |
| Disallowed (%) | 0.08 | — |

Previous structural and functional analyses of RmtC and other m^7^G1405 methyltransferases suggested that the two N-terminal subdomains of these enzymes (N1 and N2) make critical interactions in the process of 30S subunit substrate recognition (21, 22, 24). The structure of the 30S-RmtC complex reveals the molecular basis for this dependence on the N-terminal subdomains, with N1 directly contacting the 16S rRNA surface formed by helices h27, h44, and h45 (**Figure 1D**,**E**). The region of N2 closest to N1 makes additional interactions exclusively with h24 (**Figure 1E**). The N2 subdomain is larger in RmtC compared to homologs RmtA, RmtB, and ArmA (**Supplemental Figure S4**), and the map is notably weaker for this extended region. This indicates that N2 is more mobile distant from N1 and further away from the 30S surface (**Supplementary Figure S5A**), consistent with only the N2 region proximal to N1 being important for RmtC-30S subunit interaction.

Previous studies of RmtC also implicated a loop comprising residues 237 to 246 (Loop^237-246^) and adjacent C-terminal domain residues as being critical for G1405 modification, despite not directly contributing to 30S subunit binding affinity (24). The loop region was previously found to be disordered in the crystal structures of free RmtC and other G1405 rRNA methyltransferases (17, 21, 24), limiting direct insight into the function of these critical residues. In contrast, Loop^237-246^ is ordered in this 30S subunit-bound structure where these residues make direct contacts to a highly distorted region of h44 adjacent to the G1405 target nucleotide (**Figure 1F** and **Supplementary Figure S5B**).

### The RmtC N-terminal domain directs docking on a conserved 16S rRNA tertiary surface

The RmtC N-terminal domain (NTD) mediates binding and recognition of the 30S ribosomal subunit through a network of direct contacts with the 16S rRNA tertiary surface comprising h24, h27, h44, and h45 (**Figures 2** and **3**). This observation is consistent with the previously proposed overlapping interaction surface as used by the m^1^A1408 16S rRNA methyltransferases and the same requirement for mature 30S subunit as the substrate (23-25).

**Figure 2.**
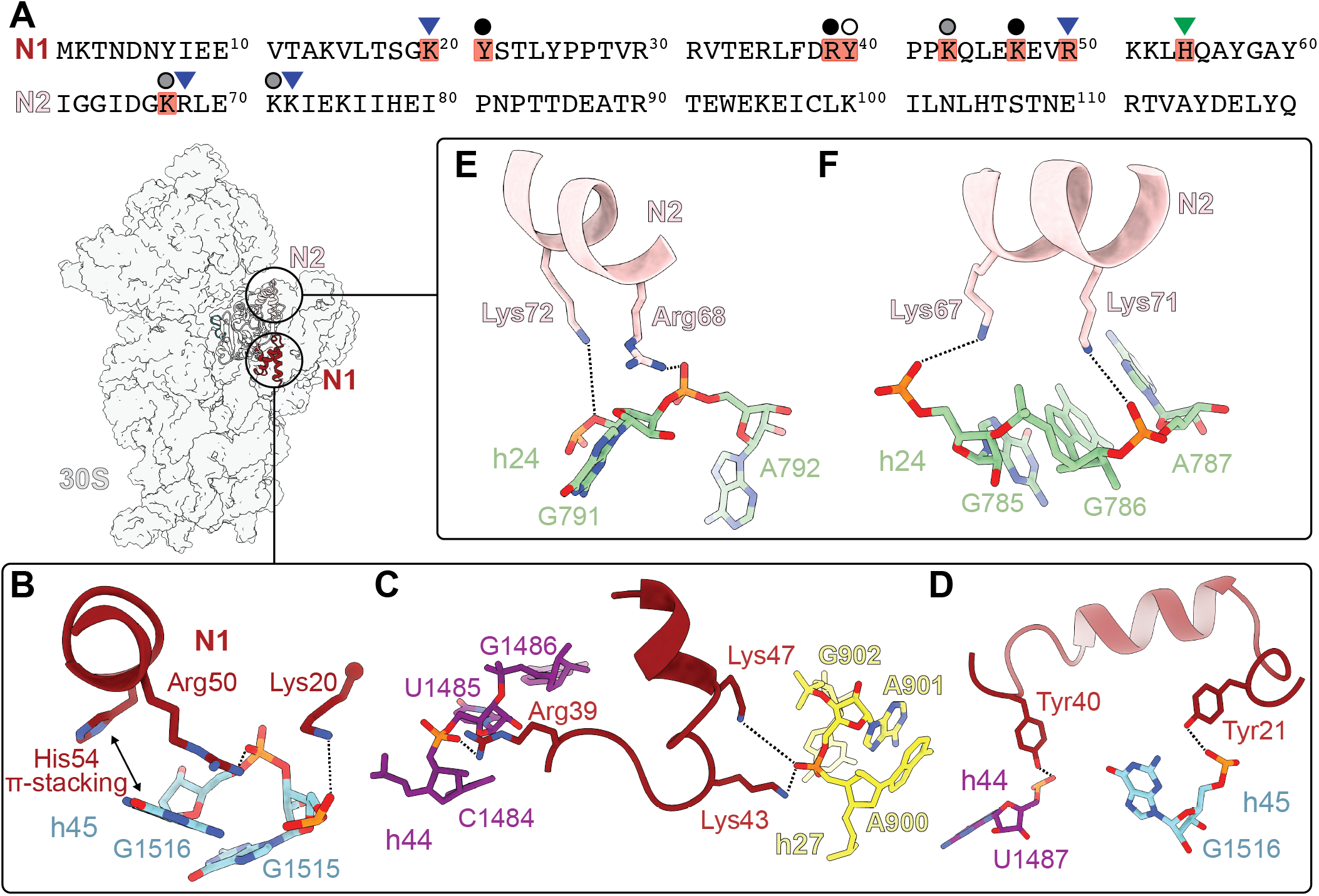
RmtC N-terminal domain residues direct recognition and binding to the 16S rRNA. **A**. Sequences of the RmtC N1 and N2 subdomains, highlighting conserved residues among m^7^G1405 methyltransferases (see **Supplementary Table S1**) observed to interact with 16S rRNA (red shading) and indicating those previously (24) found to be important for both binding and RmtC activity (blue triangle) or activity only (green triangle). New interactions identified and tested in MIC assays in the present work (see **Table 2**) are indicated with circles with shading indicating the impact of substitutions on RmtC activity: major loss (black), modest loss (gray), or no effect (white). **B**. N1 subdomain residues Lys20 and Arg50 (red) form electrostatic interactions with the phosphate backbone of h45 nucleotides G1515 and G1516 (cyan), respectively, while His54 forms a face-edge base interaction with G1516. **C**. Arg39, Lys43, and Lys47 form electrostatic interactions with U1485 (h44; purple) and A901 (h27; yellow). **D**. Tyr21 and Tyr40 (red) interact with nucleotides G1516 (h45; cyan) and G1487 (h44; purple), respectively. **E**. N2 subdomain residues Lys72 and Arg68 (pink) form electrostatic interactions with the phosphate backbone of h25 nucleotides A792 and G791 (green). **F**. Lys67 and Lys71 interact with the phosphate backbones of h24 nucleotides G785 and A787, respectively.

Two highly conserved N1 domain residues, Lys20 and Arg50, were previously shown to be essential for 30S subunit binding and m^7^G1405 modification (**Figure 2A** and **Supplementary Table S1**), through measurements of binding affinity and aminoglycoside minimum inhibitory concentrations (MIC) with RmtC protein variants (**Table 2**) (24). Our 30S-RmtC structure now reveals that Lys20 and Arg50 form critical electrostatic interactions with the phosphate backbone of h45 nucleotides G1515 and G1516, respectively (**Figure 2B**, and see **Supplementary Figure S6A**,**B** for images of map quality supporting these and the other interactions described below). Additionally, the tip of the loop between the first two N1 subdomain α-helices (residues Gly19-Lys20) is packed against the ribose of h24 residue C783 (**Supplementary Figure S7A**), thus coordinating recognition of the surface formed by 16S rRNA helices h24 and h45. Three additional conserved RmtC N1 subdomain basic residues, Arg39, Lys43 and Lys47, are also positioned to interact with h27 or h44, (**Figure 2C**,**D** and **Supplementary Table S1**). Substitution of each basic residue with glutamic acid was used to test the importance of these potential electrostatic interactions in MIC assays with kanamycin and gentamicin (**Table 2**). The R39E and K47E substitutions dramatically reduce the MIC for both aminoglycosides, indicating that these interactions with h44 and h27 are also essential for docking on the 30S subunit. In contrast, K43E substitution has a more modest impact on RmtC activity, only measurably decreasing the MIC for gentamicin, which has a lower activity in the presence of the wild-type enzyme compared to kanamycin (**Table 2**).

**Table 2.** Aminoglycoside MIC conferred by RmtC variants.

| Role/<br>Region | RmtC | Antibiotic MIC (µg/ml) |  | Reference | Figure |
| --- | --- | --- | --- | --- | --- |
|  |  | Kanamycin | Gentamicin |  |  |
| - | Wild-type | > 1024 | 1024 | - | - |
| 16S rRNA docking/<br>NTD (N1) | K20E | < 2 | < 2 | Nosrati *et al.* (24) | 2B |
|  | Y21F | 32-64 | < 2 | This study | 2D |
|  | R39E | < 2 | < 2 | This study | 2C |
|  | Y40F | >1024 | 1024 | This study | 2D |
|  | K43E | >1024 | 128-256 | This study | 2C |
|  | K47E | 8 | 2 | This study | 2C |
|  | R50E | < 2 | < 2 | Nosrati *et al.* (24) | 2B |
|  | H54A | < 2 | < 2 | Nosrati *et al.* (24) | 2B |
| 16S rRNA docking/<br>NTD (N2) | K67E | 1024 | 64 | This study | 2F |
|  | R68E | 256-512 | <2 | Nosrati *et al.* (24) | 2E |
|  | K71E | 1024 | 64 | This study | 2F |
|  | K72E | 256-1024 | 64-256 | Nosrati *et al.* (24) | 2E |
|  | K67E/K71E | 4 | <2 | This study | 2F |
|  | R68E/K72E | < 2 | <2 | Nosrati *et al.* (24) | 2E |
| h44 distortion/<br>CTD + Loop<sup>237-246</sup> | T207A | >1024 | 256 | This study | 4D |
|  | R211E | 4 | <2 | Nosrati *et al.* (24) | 4D |
|  | S239A | >1024 | 512-1024 | This study | 4C |
|  | R241E | 8 | <2 | Nosrati *et al.* (24) | 4C |
|  | M245A | <2 | <2 | Nosrati *et al.* (24) | 4D |
|  | N248A | >1024 | >1024 | This study | 4D |
| G1405/ NM6<br>NTD +CTD | Y60F | >1024 | 512-1024 | This study | 4F |
|  | Y60A | 4 | <2 | This study | 4F |
|  | K204A | 4 | <2 | This study | 4F |
|  | K236E | 8 | <2 | Nosrati *et al.* (24) | 4F |

Two tyrosine residues, Tyr21 and Tyr40, are also positioned to make hydrogen bonding interactions with h45 and h44, respectively (**Figure 2D**). The impact on RmtC activity of individual substitutions of these residues (to phenylalanine) was tested via MIC measurement, as before. Consistent with the near universal conservation of Tyr21 among all m^7^G1405 methyltransferases (**Supplementary Table S1**), the Y21F substitution dramatically reduces the MICs for both kanamycin and gentamicin (**Table 2**). In contrast, Tyr40 is less conserved across the enzyme family and the Y40F substitution has no impact on the measured MIC compared to the wild-type enzyme for either antibiotic (**Table 2** and **Supplementary Table S1**), suggesting that this hydrogen bonding interaction is not essential for RmtC binding to the 30S subunit. However, this residue is conserved as aromatic (Tyr, His or Phe) in all pathogen-associated m^7^G1405 methyltransferases, suggesting some other role for this residue that requires maintenance of its aromatic nature, e.g., in protein folding or stability, that is not apparent from the structures of the free or 30S-bound enzymes.

The RmtC N2 subdomain also contains several weakly or non-conserved residues that directly contact 16S rRNA, with Lys67, Arg68, Lys71 and Lys72 positioned to make electrostatic interactions with h24 (**Figure 2E**,**F, Supplementary Figure S6C**, and **Supplementary Table S1**). As for Arg68 and Lys72 which were previously established to contribute to RmtC binding to the 30S subunit (**Table 2**) (24), individual K67E or K71E substitutions result in a moderate reduction in resistance conferred by RmtC, whereas a K67E/K71E double substitution completely restores susceptibility to both aminoglycosides (**Table 2**). Collectively, these results support the combined importance of these residues in mediating RmtC binding to the 30S subunit.

The 30S-RmtC structure and functional studies thus reveal a set of N1 (Lys20, Tyr21, Arg39, Lys47, and Arg50) and N2 (Lys67, Arg68, Lys71 and Lys72) residues that make interactions critical for 30S binding, with a predominant role for those in the N1 subdomain. Most nucleotides contacted by these residues are positioned essentially identically in the free 30S subunit and 30S-RmtC structures (26), suggesting that this interaction network directs docking of pre-formed complementary surfaces on the enzyme and substrate to correctly position RmtC adjacent to its G1405 target site in h44. The one exception is for Arg50 where, upon RmtC binding, G1516 moves by ∽3.8 Å away from RmtC and distorts the h45 backbone (26). This movement is necessary to avoid a clash with the near universally conserved N1 subdomain residue His54 (**Figure 2B** and **Supplemental Figure S7B**), which was previously found to be essential for RmtC activity (via MIC measurements) but does not contribute measurably to RmtC-30S subunit binding affinity (24). Thus, in contrast to the other critical NTD residues, His54, together with a set of conserved C-terminal domain (CTD) residues (see below), may be essential for stabilizing functionally critical 16S rRNA conformational changes that allow RmtC to access and modify the G1405 target site.

### Loop^237-246^ and adjacent CTD residues distort h44 to induce G1405 base flipping for modification

G1405 is located in the ribosome decoding center at the top of h44 where it is Watson-Crick base paired with nucleotide C1496 and largely inaccessible in the free 30S subunit structure (26). Specifically, the G1405 N7 position is buried deep within the helix major groove and the base edge is fully occluded by h45 nucleotides G1517 and A1518 (**Supplemental Figure S8**). Thus, significant distortion of h44 surrounding the RmtC binding site is required for the enzyme to access the target site on G1405. To accomplish this, RmtC flanks h44 with its N1 domain positioned against the h44 minor groove surface one helical turn below G1405, while Loop^237-246^ contacts the same groove directly opposite G1405. Between these two regions, part of the loop connecting core β-strands β4 and β5 (residues Arg211 to Glu214) is positioned over the intervening major groove surface. Collectively, the interactions mediated by these regions of RmtC induce a ∽18 Å movement of G1405 from its original position in the free 30S subunit (**Figure 3A**), abolishing its base pairing with C1496, and flipping the nucleotide into the enzyme active site. The remainder of h44 (from C1412 to G1488) remains unaltered (**Figure 3B**).

Adjacent to G1405, nucleotides U1406, C1407, A1408 and C1409 are typically base stacked and interact with nucleotides on the complementary strand of h44. In contrast, when RmtC is bound, these nucleotides are distorted to accommodate the enzyme, with Loop^237-246^ interacting with the h44 minor groove surface, opposite the flipped G1405 (**Figure 4** and **Supplementary Figure S6D**). Through interaction with RmtC residues Ser239 and Arg241, nucleotide U1495 is moved 3.7 Å and its base rotated ∽145° towards RmtC (**Figure 4C**). Of these interactions, the most critical appears to be the near universally conserved Arg241 sidechain with the U1495 phosphate backbone as an R241E substitution was previously found to abolish RmtC activity in the MIC assay (24) (**Table 2** and **Supplementary Table S1**). In contrast, a S239A substitution results in a small reduction in MIC for gentamicin only, consistent with weaker conservation only in acquired enzyme at this site, with glycine most predominant including for all drug-producer (intrinsic) homologs.

**Figure 3.**
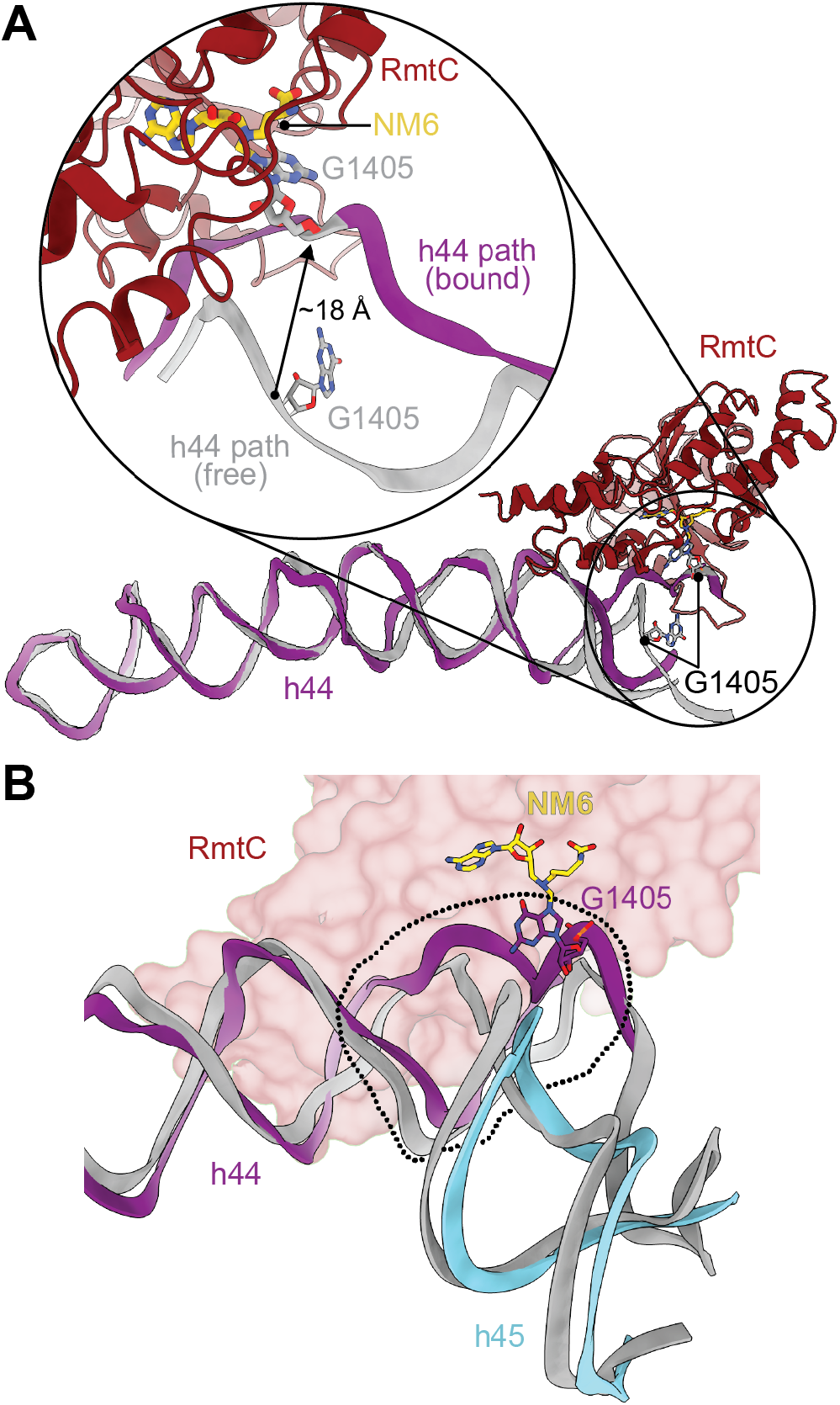
RmtC binding induces distortion of h44 surrounding the G1405 target nucleotide. **A**. RmtC (red) binds to h44 surrounding G1405, inducing a disruption of the h44 rRNA path that results in G1405 moving ∽18 Å and with a rotation around its phosphodiester backbone to flip the nucleobase into the enzyme active site for modification. The region of h44 distal from RmtC does not significantly differ in structure from the free mature 30S subunit (PDB ID 7OE1). **B**. Only the region of h44 proximal to G1405 (within the dotted outline) is distorted upon RmtC binding; the remainder of h44 (purple) and h45 (cyan) are essentially unchanged compared to the free mature 30S subunit.

**Figure 4.**
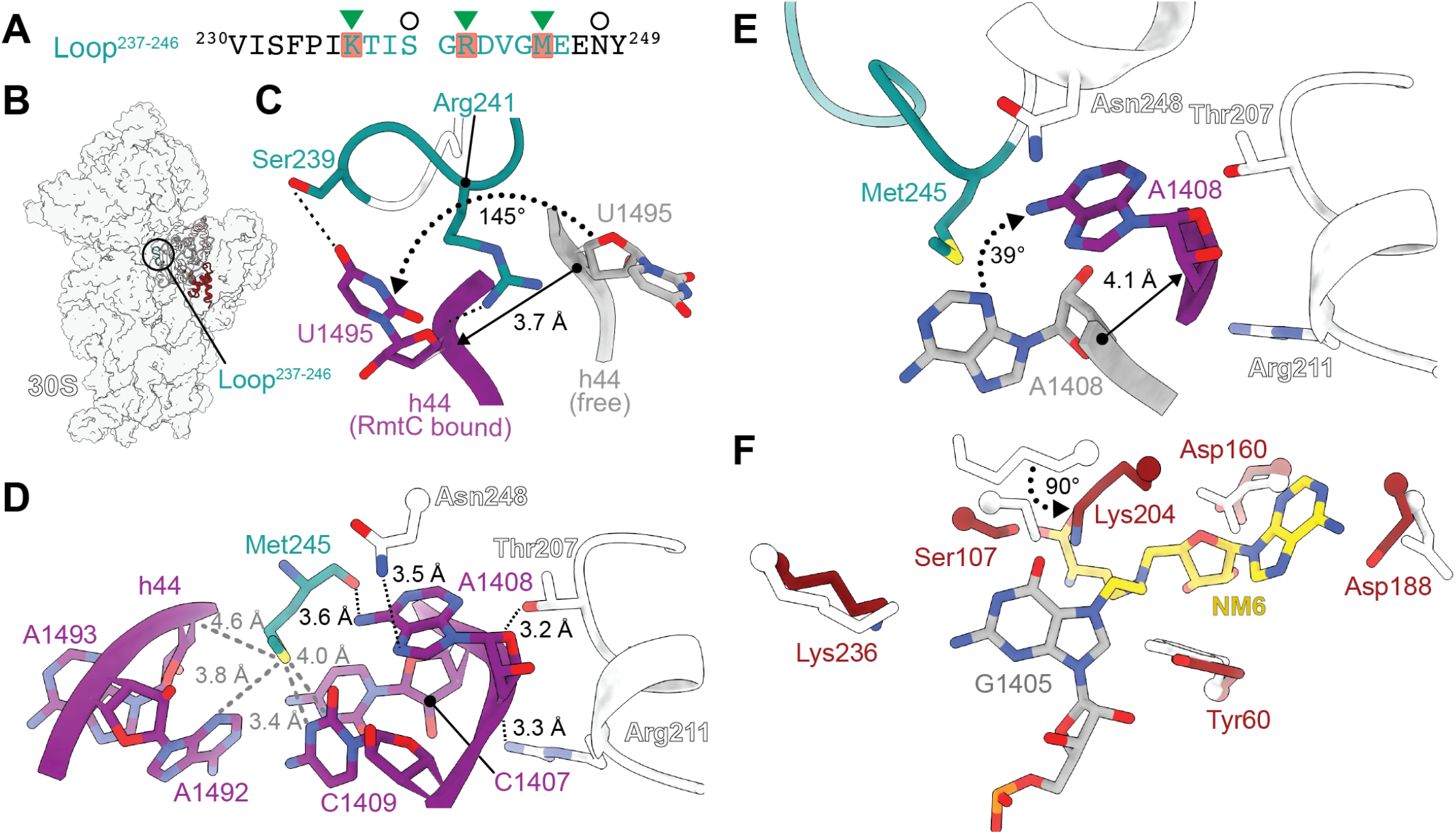
RmtC Loop^237-246^ and adjacent CTD residues direct functionally essential alterations in h44 structure. **A**. Sequence of RmtC Loop^237-246^, highlighting conserved residues among m^7^G1405 methyltransferases (see **Supplementary Table S1**) observed to interact with 16S rRNA (red shading) and indicating those previously found (24) to be important for RmtC activity (green triangle). New interactions identified and tested in MIC assays in the present work (see **Table 2**) are indicated with circles. **B**. Overview of the 30S-RmtC complex with Loop^237-246^ (teal) circled. **C**. Interaction of RmtC residues Ser239 and Arg241 promote movement and flipping of nucleotide U1495, to accommodate the distorted position of G1405 required for modification. **D**. The Met245 sidechain (teal) forms a network of van der Waals interactions with a h44 rRNA pocket comprising C1407, A1408, C1409, A1492, and A1493 (purple). The Met245 backbone and Thr208, Arg211, and Asn248 side chains also coordinate A1408 with multiple interactions. Thr207 and Asn248 (white) form interactions with h44 nucleotide A1408 (purple), while Arg211 (white) contacts the phosphate backbone. Note that Thr208, Arg211, and Asn248 are outside of Loop^237-246^ but are located in the CTD and close in the RmtC structure. **E**. Changes in h44 structure around A1408 promoted by RmtC residues Met245, Thr207, Arg211, and Asn248. **F**. Residues in the 30S-RmtC complex structure (red) are largely found in the same orientation as in the free RmtC-SAH complex (white; PDB ID 6PQB). Only Lys204 adopts a significantly altered position when RmtC binds to the 30S subunit, with the sidechain rotated ∽90° and oriented towards to the G1405 (grey) base N7 modification site.

Four other residues make additional interactions on the rRNA strand containing G1405 to further support distortion of h44: Loop^237-246^ residue Met245, and nearby C-terminal domain residues, Thr207, Arg211 and Asn248 (**Figure 4D** and **Supplementary Figure S6E**,**F**). Met245 is highly conserved among RmtC homologs (or replaced by another hydrophobic residue, leucine) and its substitution with the smaller hydrophobic residue alanine was previously shown to abolish enzyme activity (**Table 2**) (24). The Met245 side chain is positioned within van der Waals interaction distance of A1408 on the h44 minor groove surface, and the adjacent nucleotides C1407 and C1409, and A1492 and A1493 on the opposite side of h44 (**Figure 4D**). The Met245 backbone carbonyl oxygen, as well as the side chains of Thr207, Asn248, and Arg211 are also positioned to interact with the A1408 base or phosphate backbone (**Figure 4D**). Arg211 was previously identified as being important for RmtC activity, as an R211E substitution significantly reduced the conferred MIC (**Table 2**), while not contributing measurably to RmtC-30S subunit binding affinity (24). This finding can now be rationalized through the role of Arg211 in stabilizing the altered position of A1408. Although more commonly found in homologs from pathogenic bacteria, Thr207 and Asn248 are more modestly conserved (**Supplementary Table S1**). Consistent with this observation, individual T207A or N248A substitutions have only limited or no impact on the MIC, respectively, and thus do not appear individually critical for RmtC activity (**Table 2**). Collectively, however, the interactions made by these four residues result in the movement of the A1408 nucleotide backbone ∽4.1 Å inwards and rotated 39° towards RmtC and away from h44 (**Figure 4E**). As such, interactions on each side of h44 mediated by Loop^237-246^ and nearby CTD residues in combination with those made by N1 domain residues (in particular, His54) allow RmtC to act as a pincer, promoting and stabilizing a major local distortion of h44 to make G1405 accessible for methylation.

### Flipped G1405 is precisely positioned for modification by base stacking and stabilization of the modification site

As noted earlier, covalent attachment of NM6 to G1405 captured the 30S-RmtC complex in a state immediately following catalysis. Several conserved RmtC active site residues surround both NM6 and G1405, optimally positioning the guanosine base for modification at its N7 position (**Figure 4F** and **Supplementary Figures S6G-J and S9**). As previously observed in the structure of the free enzyme with SAH (24), the near universally conserved Asp160 contacts the NM6 ribose in an interaction broadly conserved in Class I methyltransferases (27), while Ser107 is positioned to interact with the NM6 carboxyl group (**Figure 4F** and **Supplementary Table S1**). Additional, potentially critical interactions supporting catalysis of modification are also now revealed in the 30S-RmtC complex structure. The side chain of Tyr60 is located between the NM6 adenosine and G1405, forming face-edge base stacking interactions with each nucleobase. Tyr60 is universally conserved as an aromatic residue (**Supplementary Table S1**) and, consistent with this limited natural variation, a Y60F substitution only very modestly decreases the enzyme activity whereas a Y60A substitution completely abolishes activity (**Table 2**). Like other residues that contact NM6, Tyr60 adopts a similar position in the 30S-RmtC complex structure as compared to the free RmtC-SAH complex (24) (**Figure 4F**), suggesting that Tyr60 coordinates the cosubstrate in a conformation primed for modification of G1405 upon 30S subunit binding.

Finally, residues Lys204 and Lys236 exclusively contact G1405 within the RmtC active site (**Figure 6**). The Lys236 sidechain is positioned towards the G1405 base, and a K236E substitution reduces the MIC of both tested aminoglycosides (**Table 2**). Substitution with alanine of the universally conserved Lys204 results in complete loss of aminoglycoside resistance (**Table 2** and **Supplementary Table S1**). Unlike the other residues contacting NM6 and G1405, the Lys204 side chain rotates ∽90° from its position in the RmtC-SAH structure (24) to interact directly with the modified N7 of G1405 and thus appears critical for positioning the base and directing its modification (**Figure 4F**). Collectively, these observations support the critical importance of Lys204 in engaging G1405 to direct catalysis of N7 modification.

## Discussion

The ribosome’s essentiality for bacterial growth and survival makes it a hub for cellular regulation and thus an important antibiotic target and subject of associated resistance mechanisms. In this work, we determined an overall 3.0 Å cryo-EM structure of the aminoglycoside-resistance 16S rRNA (m^7^G1405) methyltransferase RmtC bound to the *E. coli* 30S subunit, supported by complementary functional analyses. This work has revealed the molecular mechanism by which RmtC docks on the 30S subunit via a complex 16S rRNA tertiary surface and generates significant local distortion of h44 to reposition G1405 in the enzyme active site for modification.

RmtC binding to the 30S subunit is primarily directed by N1 and N2 subdomain residues which form interactions with the 16S rRNA sugar-phosphate backbone across a complex RNA tertiary surface comprising h24, h27, h44 and h45. The N1 subdomain and Loop^234-246^ in the CTD contact opposite sides of h44 and act as a pincer to drive a major distortion of h44 and thus make G1405 accessible for modification. In particular, His54 in the N1 domain and three key residues in Loop^234-246^ make functionally critical interactions with h44 that do not appear to contribute to 30S subunit binding affinity, but instead act to stabilize the distorted h44 structure. The role of these essential residues may thus to be offset a substantial energetic cost associated with h44 disruption. The flipped G1405 is also coordinated by N1 subdomain residue Tyr60 and CTD residues Lys204 and Lys236 to precisely position the nucleobase for modification. While target nucleotide base flipping is a common strategy employed by diverse DNA and RNA methyltransferases (23, 28, 29), the occluded location of G1405 in h44 of mature 30S subunits makes base flipping a prerequisite for G1405 modification. RmtC thus appears to possess two groups of residues important for substrate recognition and enzyme activity: one primarily in the N1 subdomain that directs initial docking on the 30S subunit and a second, surrounding the opening to the SAM binding pocket, that directs h44 distortion and positioning of G1405 for methylation (**Figure 5A**,**B**). This mechanism is distinct from other 16S rRNA methyltransferases that also recognize h44.

**Figure 5.**
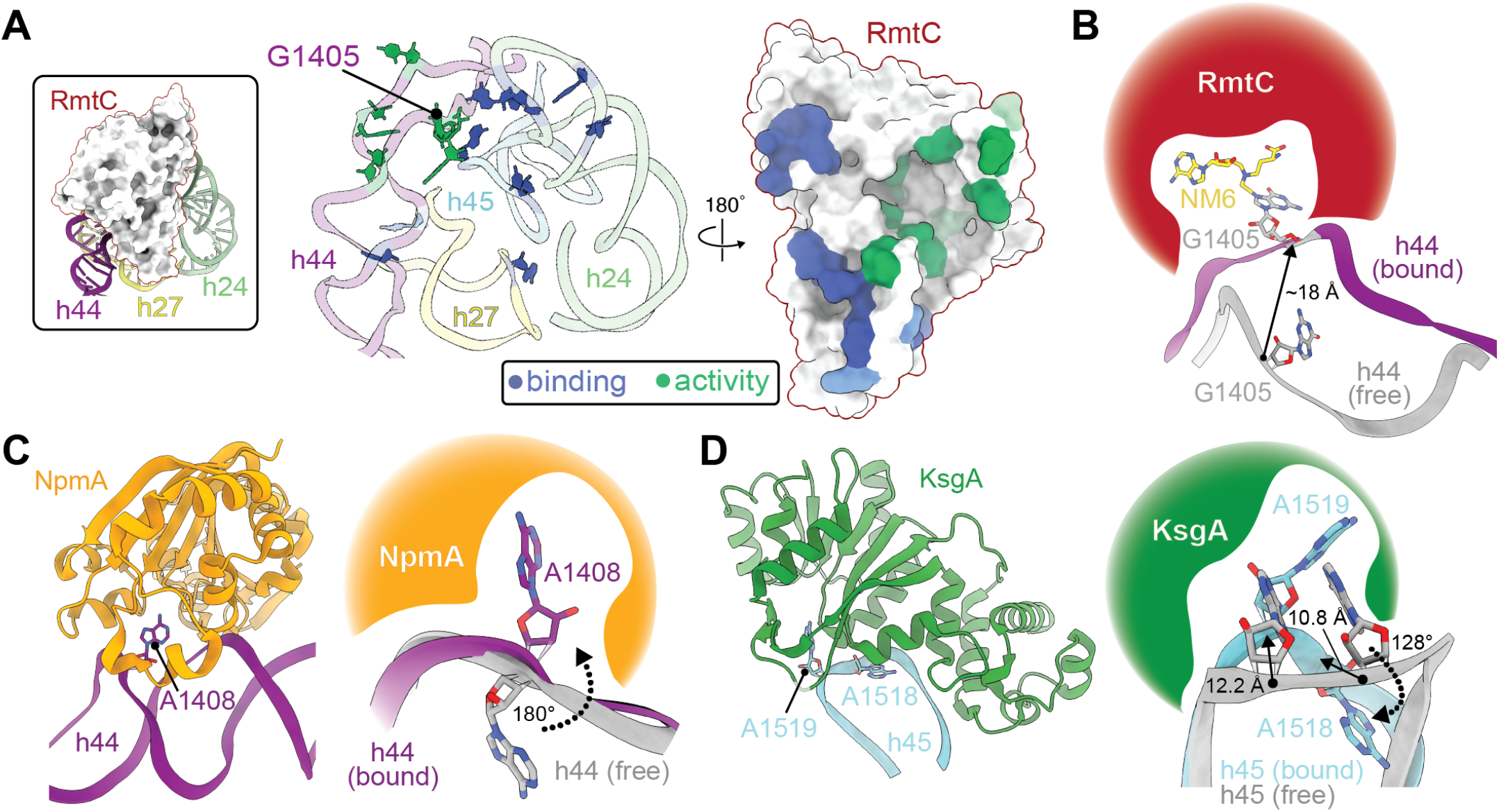
Mechanism of substrate recognition and modification by RmtC and other structurally characterized 16S rRNA methyltransferases. **A**. Views of the 16S rRNA and RmtC interaction interface surfaces highlighting locations of contacts important for both enzyme-substrate binding affinity and thus enzyme activity (blue), or enzyme activity only (green). **B**. Major distortion of h44 is required to flip G1405 into the RmtC active site. **C**. Overview of NpmA (orange) interaction with h44 (*left*; PDB ID 4OX9) and zoomed view of the limited local distortion at A1408 necessary to flip the target nucleotide into the NpmA active site (*right*). **D**. Overview of KsgA (green) interaction with h45 (*left*; PDB ID 7O5H) and zoomed view of the distortion at A1518/ A1519 necessary to flip the target nucleotide (A1519) into the KsgA active site.

Prior to the current work, only two structures of rRNA methyltransferases bound to the 30S subunit have been determined: the aminoglycoside-resistance 16S rRNA (m^1^A1408) methyltransferase NpmA (23), and the m_2_^6,6^A1518/m_2_^6,6^A1519 dimethyltransferase KsgA, which is involved in 30S biogenesis (30, 31). Like RmtC, both NpmA and KsgA contact the complex 16S rRNA surface formed by h24, h27, h44 and h45, and induce flipping of their target nucleotide for modification (**Figure 5C**,**D**) (23, 30, 31). Although the two aminoglycoside-resistance enzymes NpmA and RmtC recognize the same conserved 16S rRNA tertiary surface (23), there are several differences in how this is accomplished. NpmA docking on 30S is mediated by three regions within the Class I methyltransferase core fold (β2/3-, β5/6-, and β6/7-linkers) rather than an extended N-terminal domain, as for RmtC. The structure of the NpmA bound to the 30S subunit also revealed A1408 to be flipped from h44 via a highly localized ∽180° rotation of the 16S rRNA backbone at this nucleotide, with essentially no other disruption of the 16S rRNA structure (23) (**Figure 5C**). Finally, NpmA stabilizes the local helical distortion at A1408 using a single arginine residue (Arg207) contact to the phosphate group that is critical for enzyme activity but does not contribute to 30S binding affinity (23, 25); the flipped conformation is further stabilized by π-stacking of the A1408 base between two universally conserved tryptophan residues. In contrast, likely due to the greater distortion required to flip G1405 from h44, RmtC relies on a more extensive interaction network to facilitate h44 distortion and uses distinct contacts by two key residues (Tyr60 and Lys204) to position the base for methylation. For KsgA to access and modify A1518 and A1519, a distortion of the surrounding highly conserved h45 tetraloop nucleotides (G1516 to A1519) is required (31, 32) (**Figure 5D**). In the structural snapshots captured of the 30S-KsgA complex, the phosphate backbone of A1519 moves ∽11-12 Å into the KsgA active site, while A1518 rotates ∽128° away from the enzyme. Methylation of A1518 is proposed to follow a similar movement into KsgA with A1519 rotating away following its modification (31, 32). As for RmtC and NpmA, these movements appear essential for proper positioning of the nucleotides to be methylated. In summary, while all three enzymes exploit similar conserved 30S subunit features for specific substrate recognition, the mechanisms by which this is accomplished and the extent of necessary conformational change in the rRNA are highly unique for each.

In addition to being the target of the aminoglycoside-resistance methyltransferases, the 16S rRNA of the ribosomal decoding center is decorated with other modifications incorporated by four housekeeping (intrinsic) methyltransferases, RmsH (m^4^C1402), RsmI (Cm1402), RsmF (m^5^C1407), and RsmE (m^3^U1498). Like RmtC, NpmA and KsgA, these enzymes require mature 30S subunit as their substrate and thus likely exploit the same 16S rRNA tertiary surface as a key component of their substrate recognition mechanism. Interestingly, in contrast, 16S rRNA modifications more distant from the decoding center, as well as both antibiotic-resistance and intrinsic modifications of the 23S rRNA are incorporated by enzymes that act on intermediates during the process of ribosome subunit assembly. The overlapping sites of the aminoglycoside-resistance and intrinsic decoding center 16S rRNA methyltransferases also results in potential for prior modifications to influence subsequent substrate recognition and modification. Indeed, some, albeit conflicting, evidence suggests aminoglycoside-resistance modifications can alter intrinsic modification levels and thus impact translation and bacterial fitness (33-35). However, further careful analyses are needed to mechanistically define such competition between 16S rRNA modifications and the resulting impact on ribosome function.

In conclusion, our structure and complementary functional analyses provide critical new insight on 30S subunit recognition and aminoglycoside-resistance modification by RmtC. In addition to already well-established mechanisms of clinical aminoglycoside resistance via drug modification or efflux, the increasing global dissemination and prevalence of these rRNA modification enzymes (including ArmA and RmtA-H), represents a major additional threat to aminoglycoside efficacy, including the latest generation drugs (4, 15, 36). The high functional conservation identified here of most essential RmtC residues suggests that all m^7^G1405 methyltransferases–whether of pathogen or drug-producer origin–likely rely on the same extensive interaction networks for 30S binding and h44 reorganization, and thus mediate modification activity through a conserved molecular mechanism. These insights therefore provide a firm foundation for future work to counter the action of RmtC and its homologs to prevent aminoglycoside resistance via rRNA modification.

## Materials and Methods

### 30S-RmtC specimen preparation

RmtC (UniProt code Q33DX5) was expressed in *E. coli* using a modified pET44 plasmid encoding a synthetic *E. coli* codon-optimized gene (“pET44-RmtC”; GenScript) and subsequently purified by Ni^2+^-affinity and gel filtration chromatographies, as described previously (24). Purified protein was concentrated to ∼1 mg/ml and flash frozen for storage at -80 °C before use. *E. coli* (MRE600) 30S ribosomal subunits and SAM analog NM6 [5′-(diaminobutyric acid)-N-iodoethyl-5′-deoxyadenosine ammoniumhydrochloride] were prepared as previously described (25, 29). A mixture of RmtC (5 μM), *E. coli* 30S subunit (1.5 μM), and NM6 (20 μM) was prepared and 3.0 μl applied to freshly glow-discharged grids (1.2/1.3 300 mesh Cu Quantifoil), with blotting for 3 s at >90% humidity before freezing in liquid ethane using a CP3 plunger (Gatan). Grids were stored in liquid nitrogen until used for data collection.

### Electron microscopy, image processing and data analysis

Data were collected at the Pacific Northwest Cryo-EM Center (PNCC) on a Titan Krios microscope (FEI) operating at 300 keV with a K3 direct electron detector (Gatan). A total of 4837 micrographs were collected using a defocus range of −0.5 to −3.5 μm at a 29,000× magnification in super-resolution mode (2×-binned with a 0.7983 Å/pixel size). Micrographs were collected as 50 frames with total 49 – 51 e^-^/Å^2^ dose over 2.46 s exposure.

Image processing was conducted in Relion 3.1 (37). Motion correction and dose weighing was conducted with MotionCorr2 (38), and contrast transfer function parameters estimated by Gctf (39). An unpublished *E. coli* 30S cryo-EM map was used for 3D-reference based autopicking. Particles were 4×-binned before further processing. Incorrectly selected particles were discarded after reference-free 2D class averaging. An *ab initio* model with C1 symmetry was created and used as a reference map for 3D refinement. Classification (3D) without alignment was performed to discard non-30S subunit particles (**Supplemental Figure S2**), followed by focused classification using an RmtC mask. Another round of three-dimension classification without alignment was performed to discard 30S particles without RmtC. Final classes containing the 30S-RmtC complex were assessed for any conformational variation, combined to maximize the number of RmtC orientations in the final reconstruction, unbinned, and subject to 3D refinement, particle polishing, CTF refinements, and post-processing, to yield a final overall 3.0 Å reconstruction, as calculated from Fourier shell correlations (FSC) at 0.143 (**Supplemental Figures S3C**). Analysis of the angular distribution of particles comprising the final map indicates some preferred orientation in the dataset, but still exhibits complete coverage (**Supplemental Figure S3E**). Local resolution was calculated in Relion (37), with a range from 2.8 to 8.0 Å (**Supplemental Figure S3F**).

In addition to the final processing of the complete 30S-RmtC map, two distinct rounds of multibody refinement were performed on the particles to assess the potential for improvement in the reconstruction of each defined body (40). Based knowledge that individual masked entities may act as a rigid body but differ in relative orientation to each other due to differences in 30S subunit conformation (41, 42), separate masks were used for the RmtC, 30S head, and 30S body, or the 30S and RmtC-h44 subcomplex. Refined maps were post-processed in both Relion (37) and in Phenix (43), and the maps with the best density in each region were used for model building.

An initial coordinate file model was created by docking the *E. coli* 30S subunit after *de novo* modeling of the N-mustard 6 (NM6)-modified G1405 (SMILES: O=C1C(N(CCN(CCC([NH3+])C([O-])=O)CC2C(O)C(O)C(N3C(N=CN=C4N)=C4N=C3)O2)CN5C6OC(COP([O-])=O)C(O)C6O)=C5N=C(N)N1) into the map using Coot and Phenix (44, 45), based on map density and the previously solved structure of RmtC bound to SAH. A complete RmtC model was generated with Alphafold2 (46, 47). The resulting model was real-space refined using the post-processed and individual multibody refinement maps in Phenix (43). Additional model building was conducted in Coot (45), with the previously solved RmtC-SAH structure guiding rotamer orientation for areas with poor density. The final model was validated in Phenix (43). Structure images were created using UCSF Chimera and ChimeraX (48, 49). Coordinates and all maps used for building (including the final Relion postprocessed map, Phenix sharpened map, two 3D-refined half maps, postprocessed multibody refinement maps of RmtC/30S head/30S body and 30S/RmtC-h44) were deposited in the RCSB PDB (8GHU) and EMDB (40051).

### Phylogenetic analysis and residue conservation

16S rRNA (m^7^G1405) methyltransferase sequences were retrieved from multiple BLAST searches using RmtC (Q33DX5), Sgm (Q7M0R2) and RmtB (Q76G15) as the query sequence. Included sequences were at least >25% in sequence identity and with >80% coverage to RmtC. Sequence redundancy was removed at 99% sequence similarity cutoff using Decrease Redundancy on the Expasy server. This process resulted in a set of 68 representative sequences, including 11 from drug-producing bacteria and 18 acquired by pathogenic bacteria. This sequence set was aligned using Geneious following optimization of gap opening and extension penalty (to 10 and 3 respectively). The evolutionary history was inferred using the Minimum Evolution (ME) method implemented in MEGA X (50). The evolutionary distances were computed using the Poisson correction method and are in the units of the number of amino acid substitutions per site. The bootstrap consensus tree inferred from 500 replicates and the residue propensities were calculated using Geneious.

### Minimum inhibitory concentration assays

RmtC variants (**Table 2**) were prepared in the pET-RmtC plasmid using whole-plasmid PCR approaches and confirmed by commercial automated DNA sequencing. Fresh lysogeny broth supplemented with 100 μg/ml ampicillin was inoculated with overnight *E. coli* BL21(DE3) culture containing plasmid encoding the wild-type or mutant RmtC sequence. Cells were grown to an OD_600_ of ∽0.1 at 37 °C and cells from 1 ml of culture were collected by centrifugation, washed twice with 0.5 mL PBS, and resuspended in cation-adjusted Mueller–Hinton (CA-MHB) medium to an OD_600_ of 0.1 (5 × 10^7^ cfu/ml). Cells were then diluted 50-fold with CA-MHB, and 100 μl was used to inoculate (1 × 10^5^ cfu/well) an equal volume of CA-MHB with 10 μM isopropyl β-D-thiogalactopyranoside and 4-2048 μg/ml of kanamycin or gentamicin predispensed on a 96-well plate. Wells containing medium with no antibiotic or no added cells served as controls in each set of experiments. Four to six individual colonies for each RmtC protein were tested from at least two independent transformations of bacterial cells with plasmid. Plates were incubated at 37 °C with shaking and OD_600_ readings taken after 24 h. The MIC reported was defined as the lowest concentration of antibiotic that fully inhibited growth (OD_600_ of <0.05 above background).

## Acknowledgements

Thanks to Dunham and Conn lab members Drs. Ha An Nguyen and Zane Laughlin for technical support. This work was supported by the National Institutes of Health awards R01-AI088025 (GLC and CMD), T32-AI106699 (PS), and T32-GM008602 (PS), and the Burroughs Wellcome Fund Investigator in the Pathogenesis of Infectious Disease award (CMD). This study was also supported by the Robert P. Apkarian Integrated Electron Microscopy Core (IEMC) at Emory University, which is subsidized by the Emory School of Medicine and Emory College of Arts and Sciences. A portion of this research was supported by NIH grant U24GM129547 and performed at the PNCC at OHSU and accessed through EMSL (grid.436923.9), a DOE Office of Science User Facility sponsored by the Office of Biological and Environmental Research.

