## Supporting Information for "30S subunit recognition and G1405 modification by the aminoglycoside-resistance 16S ribosomal RNA methyltransferase RmtC"

**This PDF file includes:**

Figures S1 to S10
Table S1

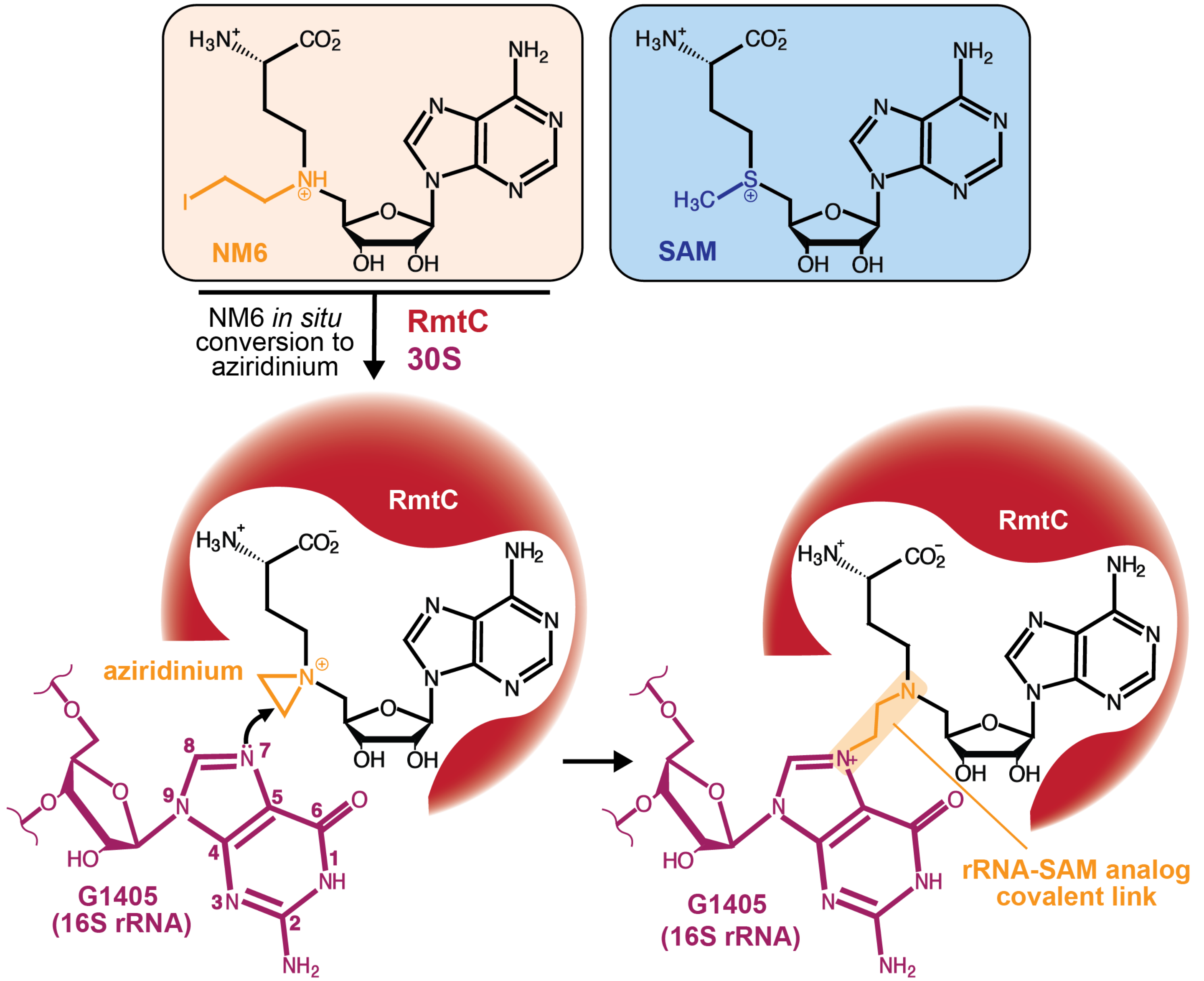


**Figure S1. Stabilization of the 30S-RmtC complex using the SAM analog NM6.** N-mustard 6 (NM6; *left box*) is an S-adenosyl-L-methionine (SAM; *right box*) analog that can be used by RmtC as a cosubstrate for 16S rRNA nucleotide G1405 (purple) modification. In the reaction catalyzed by RmtC, following *in situ* conversion to its aziridinium form, NM6 is covalently attached to the N7 nitrogen of G1405. The 30S-RmtC complex is thus captured in an immediately post-catalytic state due to RmtC's affinity for the both the 30S subunit and NM6.

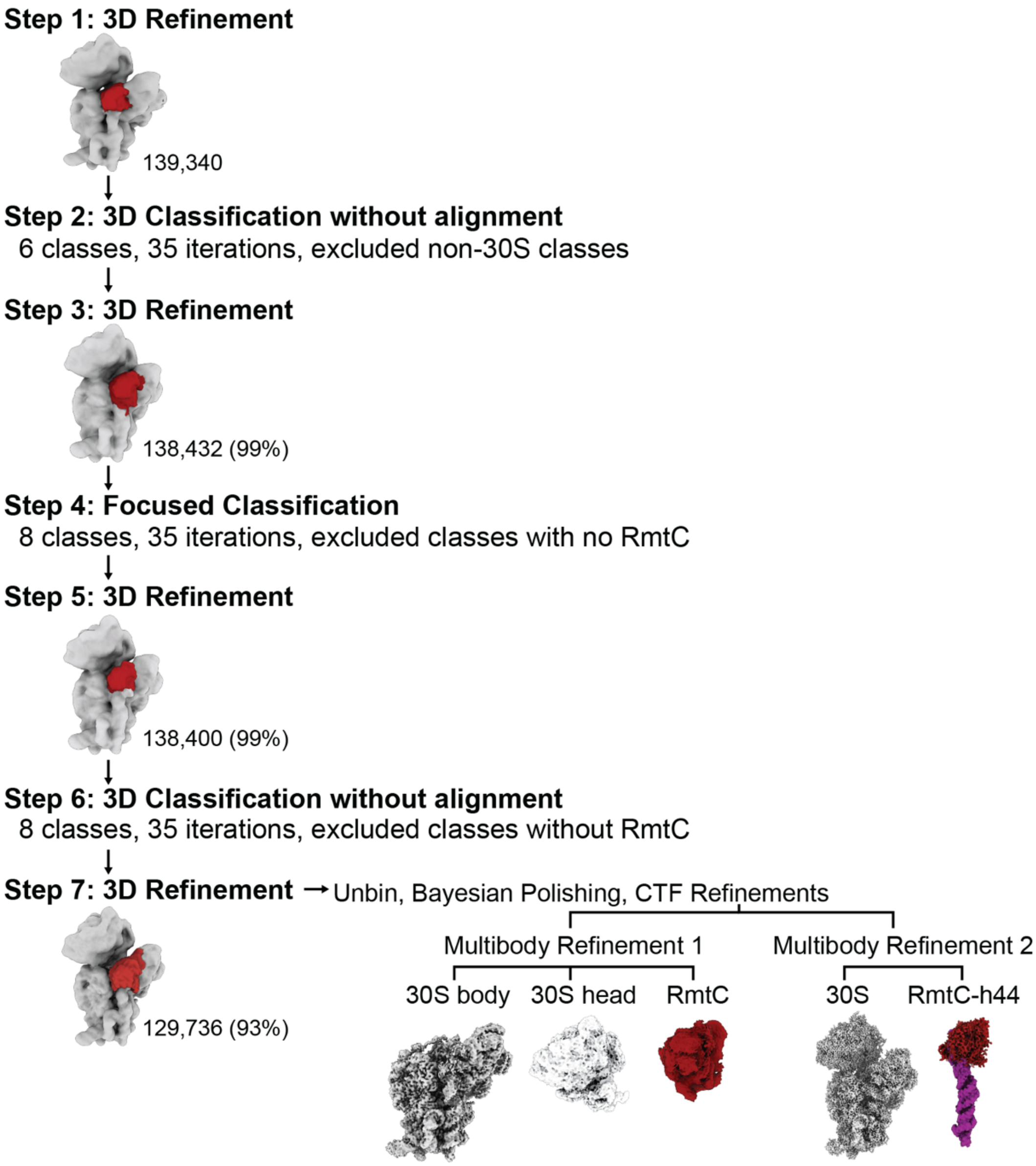


**Figure S2. Classifications of the 30S-RmtC complex.** All particles were 3D refined (**Step 1**), followed by 3D classification without refinement (**Step 2**). Classes that did not correspond to 30S subunit (dark grey) were discarded, and remaining classes were combined and subject to 3D refinement (**Step 3**). Next, a focused classification was conducted with a focused mask over the RmtC density (red; **Step 4**). Classes without RmtC (dark grey) were discarded, while classes that contained density corresponding to RmtC were combined and 3D refined (**Step 5**). This was followed by another round of 3D classification without alignment (**Step 6**), and particles without clear RmtC density (dark grey) were discarded. The remaining classes were pooled and 3D refined, and particles subsequently unbinned (**Step 7**). Unbinned particles were subject to Bayesian polishing and several rounds of refinement until resolution plateaued. Multibody refinements were performed on the final class reconstruction with masks either over the body, head, and RmtC entities, or 30S and RmtC-h44.

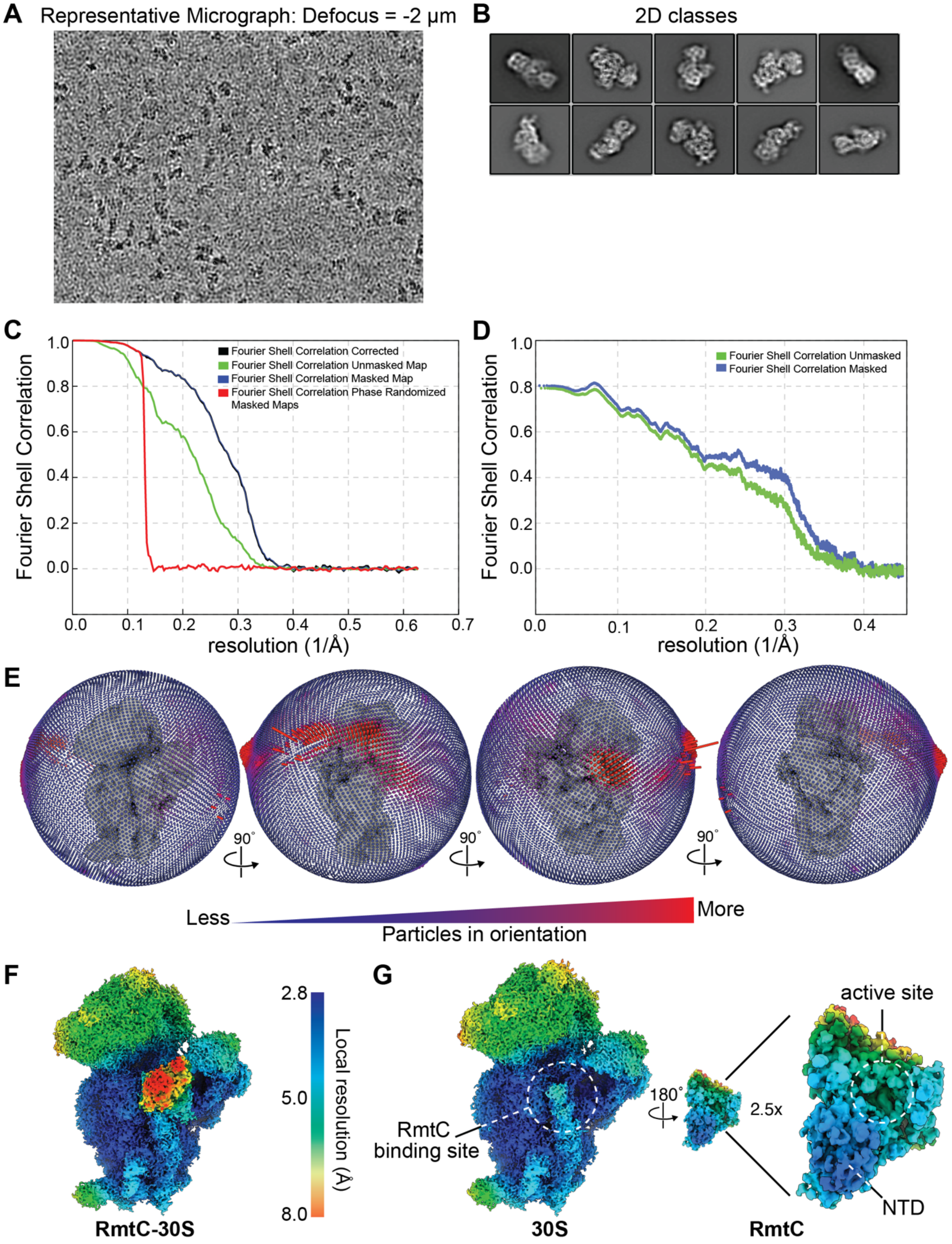


**Figure S3. Determination of cryo-EM map resolution. A**. Representative micrograph from the cryo-EM dataset, at a defocus of -2 µm. **B**. Two-dimensional class averages that were input into 3D refinement jobs. **C**. FSC plots for the 30S-RmtC final post-processed map; map resolution is 3.0 Å at $FSC_{0.143}$. **D.** Masked and unmasked map versus model FSC curve; resolution is 3.1 Å at $FSC_{0.143.}$ **E.** Angular distribution plots of particles contributing to the final map. The leftmost view is the one shown in other figures. The height (low to high) and color (blue to red) of the bars is proportional to number of particles comprising that view. **F**. Local resolution of the RmtC-30S complex using unfiltered half maps. **G**. Local resolution of the 30S with RmtC cut out and rotated 180° to show the interaction interface of each.

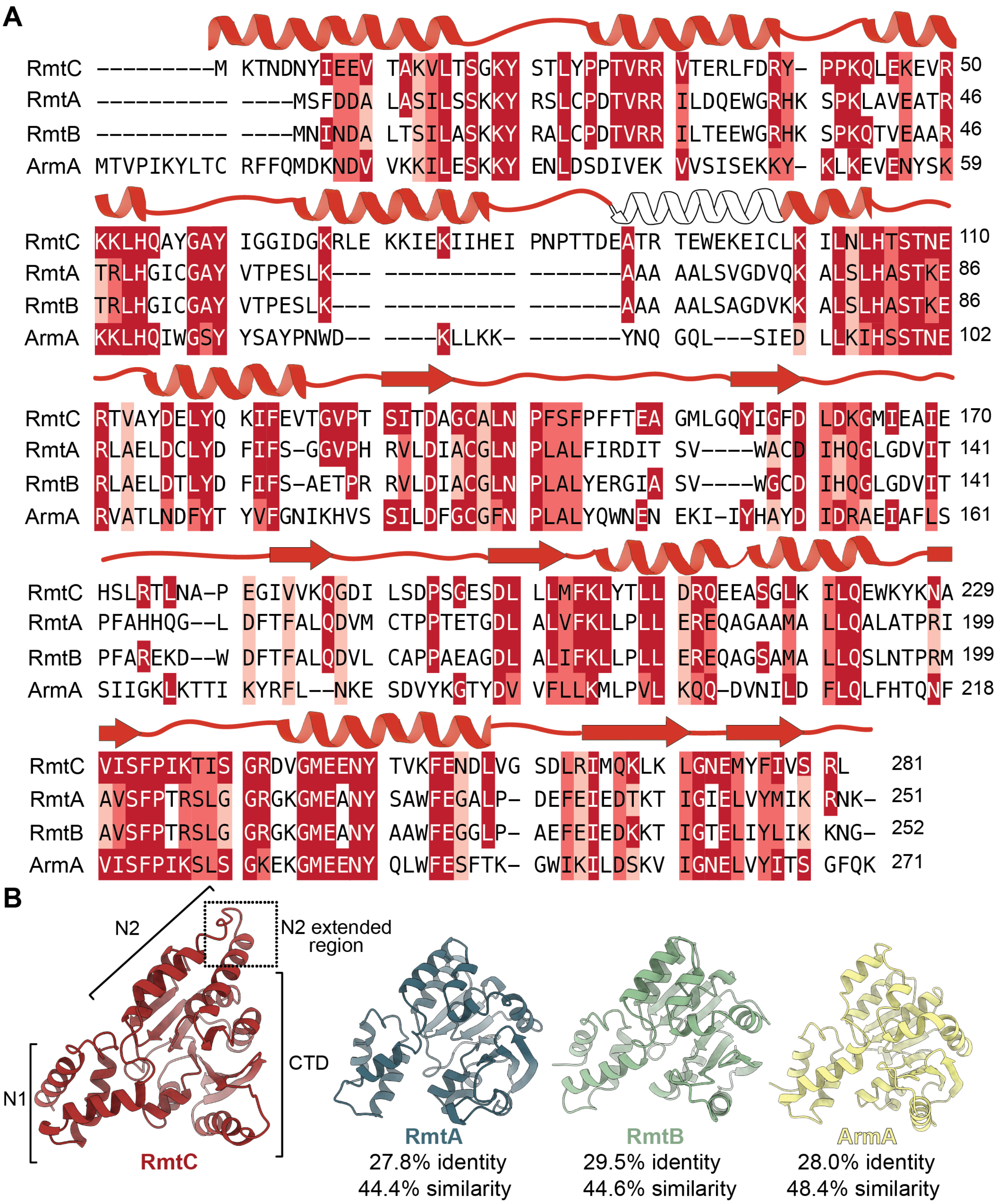


**Figure S4. Sequence and structural conservation of select m[1]G1405 methyltransferases. A**. Sequence and structural comparison of RmtC with select 16S rRNA (m[7]G1405) methyltransferase enzymes: ArmA, RmtA and RmtB. Shading indicates amino acids identical (dark red, white text) or similar to RmtC (red and light red). Protein secondary structure is shown above the corresponding sequence with conservation (red) and regions unique to RmtC (white) indicated. **B**. Structural comparison and sequence identity and similarity (determined based on the BLOSUM62 matrix) of RmtC (AF-Q33DX5-F1), RmtA (AF-Q8GRA1-F1), RmtB (PDB code 3FRI), and ArmA (AF-Q6F5A0-F1). The domains of RmtC, and the extended region of the N2 subdomain of RmtC are indicated.

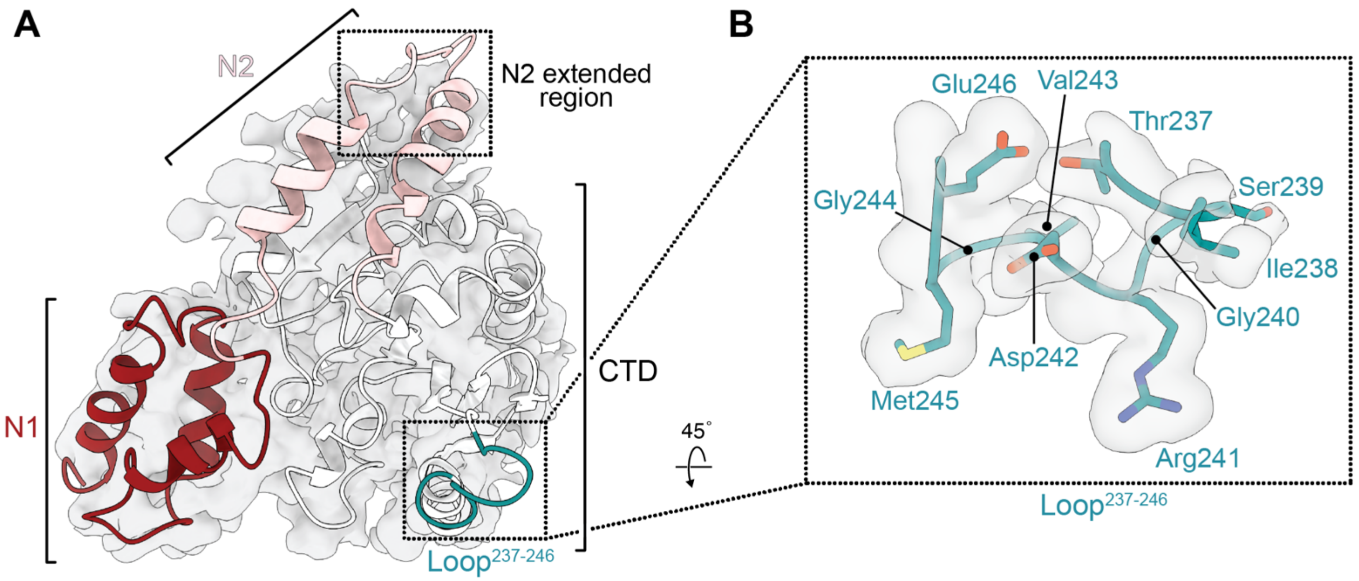


**Figure S5. Cryo-EM map for RmtC. A.** Cryo-EM map (Phenix sharpened map) corresponding to RmtC with regions of the protein indicated: N1 and N2 subdomains of the N-terminal domain (dark red and pink, respectively), C-terminal domain (white), and Loop$^{237\text{-}246}$ (teal). The N2 subdomain is extended compared to other 16S rRNA (m$^7$G1405) methyltransferases and this region is less fully resolved in the map (boxed region). **B.** Close up of the density corresponding to residues in the Loop$^{237\text{-}246}$ which becomes ordered upon 30S subunit binding (shown for the postprocessed RmtC multibody map).

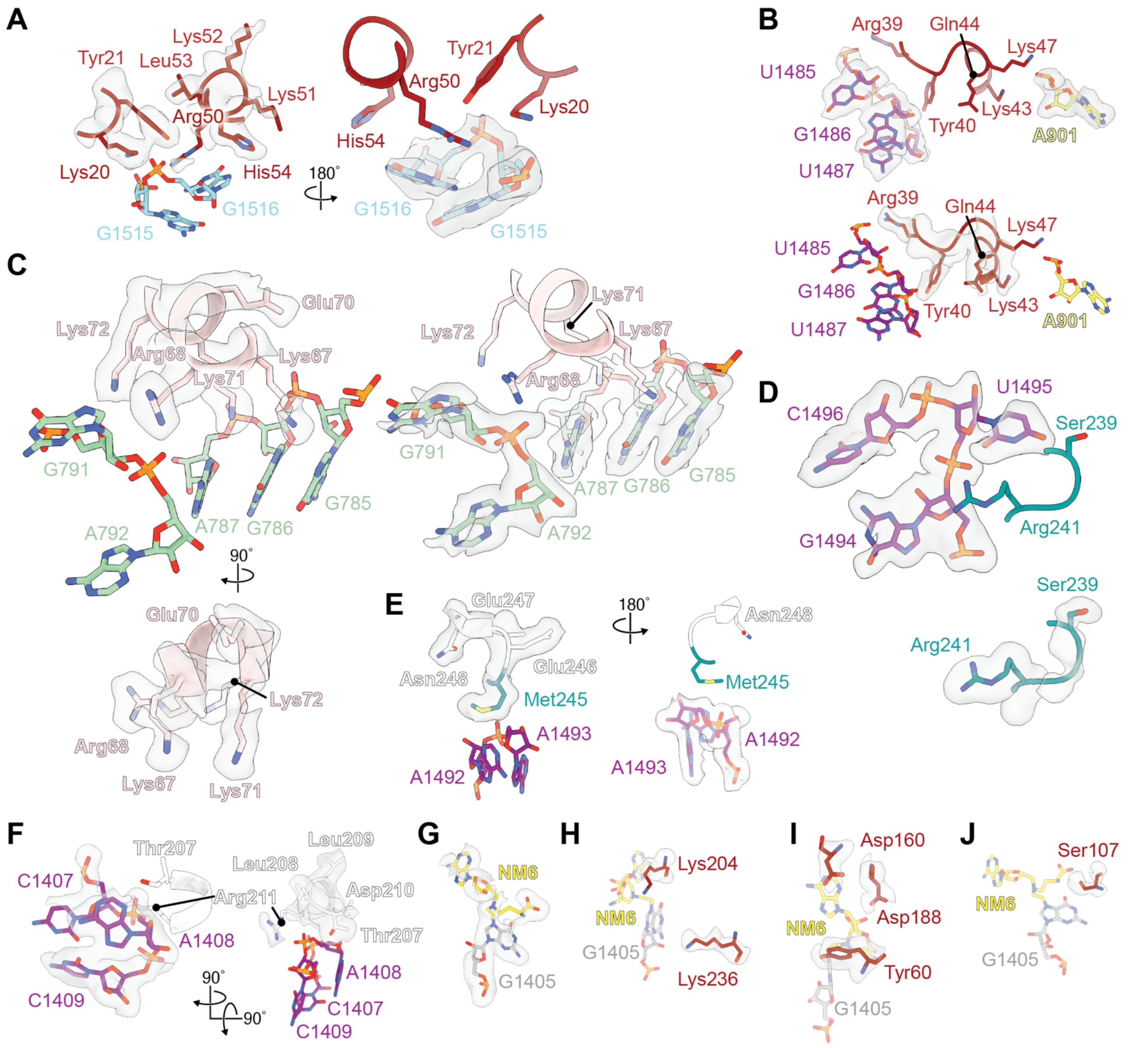


**Figure S6. Cryo-EM map quality for RmtC residues contacting 16S rRNA and NM6.** Views of the RmtC residue interactions with 16S rRNA or NM6 discussed in the main text shown with the final complete 30S-RmtC complex map (light grey). RmtC regions and rRNA helices are colored as in the main figures: RmtC N1 subdomain (dark red), N2 subdomain (pink), CTD (white) and Loop$^{237\text{-}246}$ (teal); rRNA helices h24 (green), h27 (yellow), h44 (purple) and h45 (cyan). Shown are interactions of RmtC residues: **A**. Lys20, Tyr21, Arg50 and His54 with h45 (Phenix sharpened map); **B**. Arg39, Tyr40, Lys43 and Lys47 (postprocessed RmtC-h44 multibody map) with h44 and h24 (Phenix sharpened map); **C**. Lys67, Arg68, Lys71 and Lys72 (postprocessed RmtC-h44 multibody map) with h27 (postprocessed map); **D**. Ser239 and Arg241 (postprocessed RmtC-h44 multibody map) with h44 (phenix sharpened map); **E**. Met245 and Asn248 with h44 (Phenix sharpened map); **F**. Thr207 and Arg211 with h44 (Phenix sharpened map); **G**. G1405 covalently linked to the SAM analog NM6 (postprocessed RmtC-h44 multibody map). Interaction of RmtC residues: **H**. Lys204 (postprocessed RmtC-h44 multibody map) and Lys236 (postprocessed RmtC-h44 multibody map) with G1405; **I**. Tyr60 (postprocessed RmtC-h44 multibody map), Asp160 (Phenix sharpened map), and Asp188 (postprocessed RmtC-h44 multibody map) with NM6; and, **J**. Ser107 (postprocessed map) with NM6.

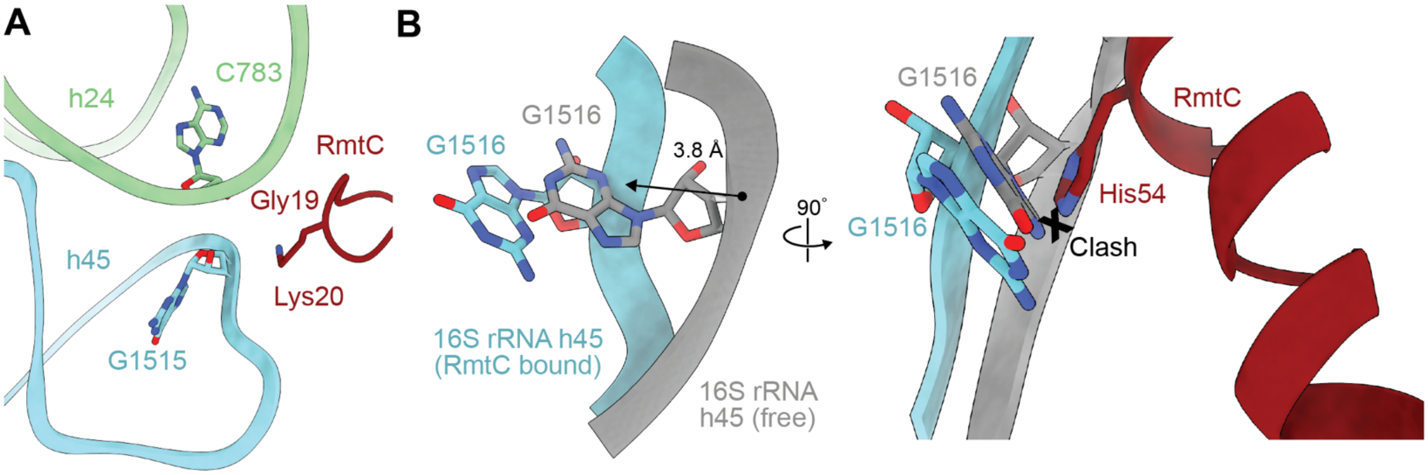


**Figure S7. N1 domain loop interactions with the h24 and h45 interface. A**. The tip of the RmtC N1 domain loop residues Gly19 and Lys20 (dark red) pack against the ribose of h24 nucleotide C783 (green), while Lys20 is also positioned to make an electrostatic interaction with the backbone of h45 nucleotide G1515. **B**. G1516 moves ~3.6 Å away from where RmtC binds compared to its position in the free mature 30S ribosome (gray; PDB ID 7OE1). This movement allows accommodation of RmtC by positioning His54 (red) to form a face-edge stacking interaction with G1516, replacing this nucleotide's normal stacking with G1515 in the free mature 30S and preventing a clash between RmtC and G1516.

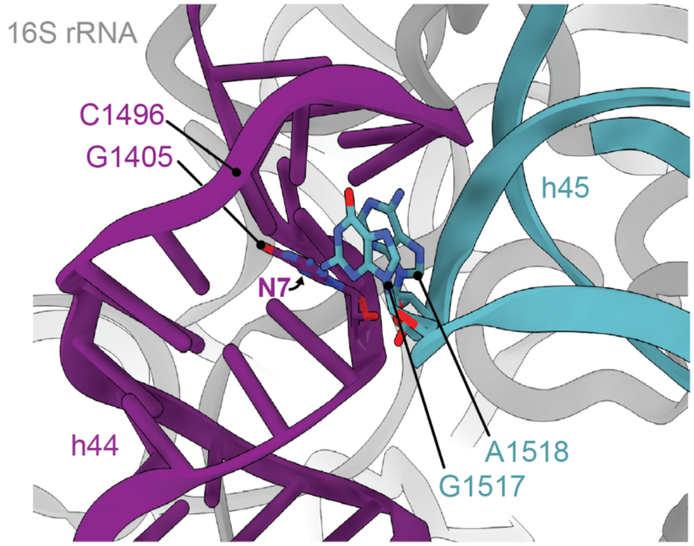


**Figure S8. The N7 modification site on G1405 is inaccessible in the free 30S subunit.** In the unbound 30S subunit (PDB ID 7OE1), nucleotide G1405 (purple) forms a Watson-Crick base pair with C1496 within h44. Additionally, nucleotides G1517 and A1518 of h45 (blue) are positioned over the h44 surface further limiting accessibility of G1405.

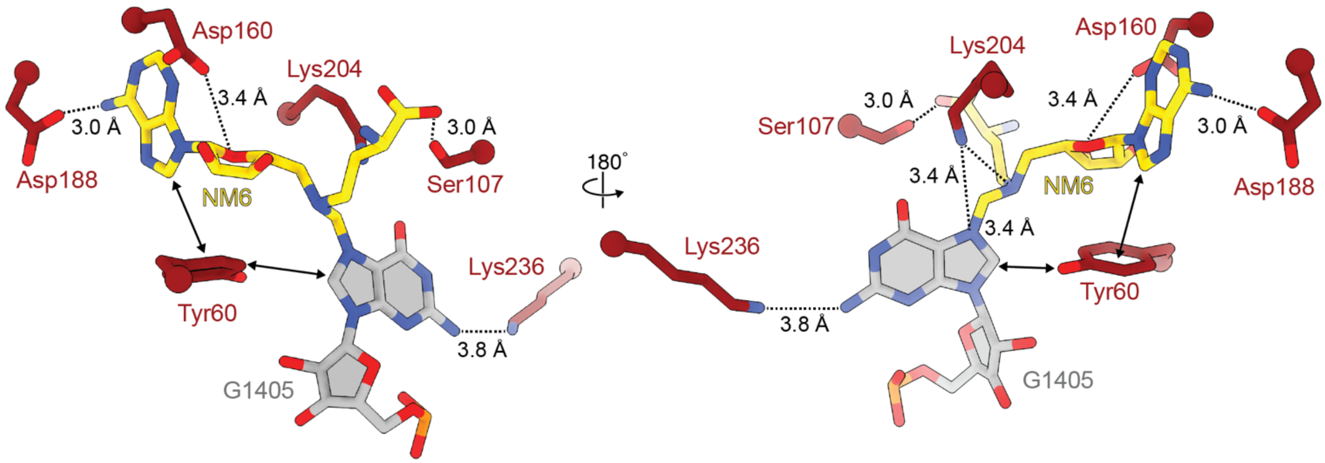


**Figure S9. Covalently attached G1405 and NM6 are surrounded by conserved RmtC residues critical for activity.** Two views of the RmtC residues (red) that coordinate the SAM analog and position G1405 (grey) for modification. Functional analyses reveal the interactions made by Tyr60 and Lys204 are especially critical for RmtC activity (see **Table 2**).

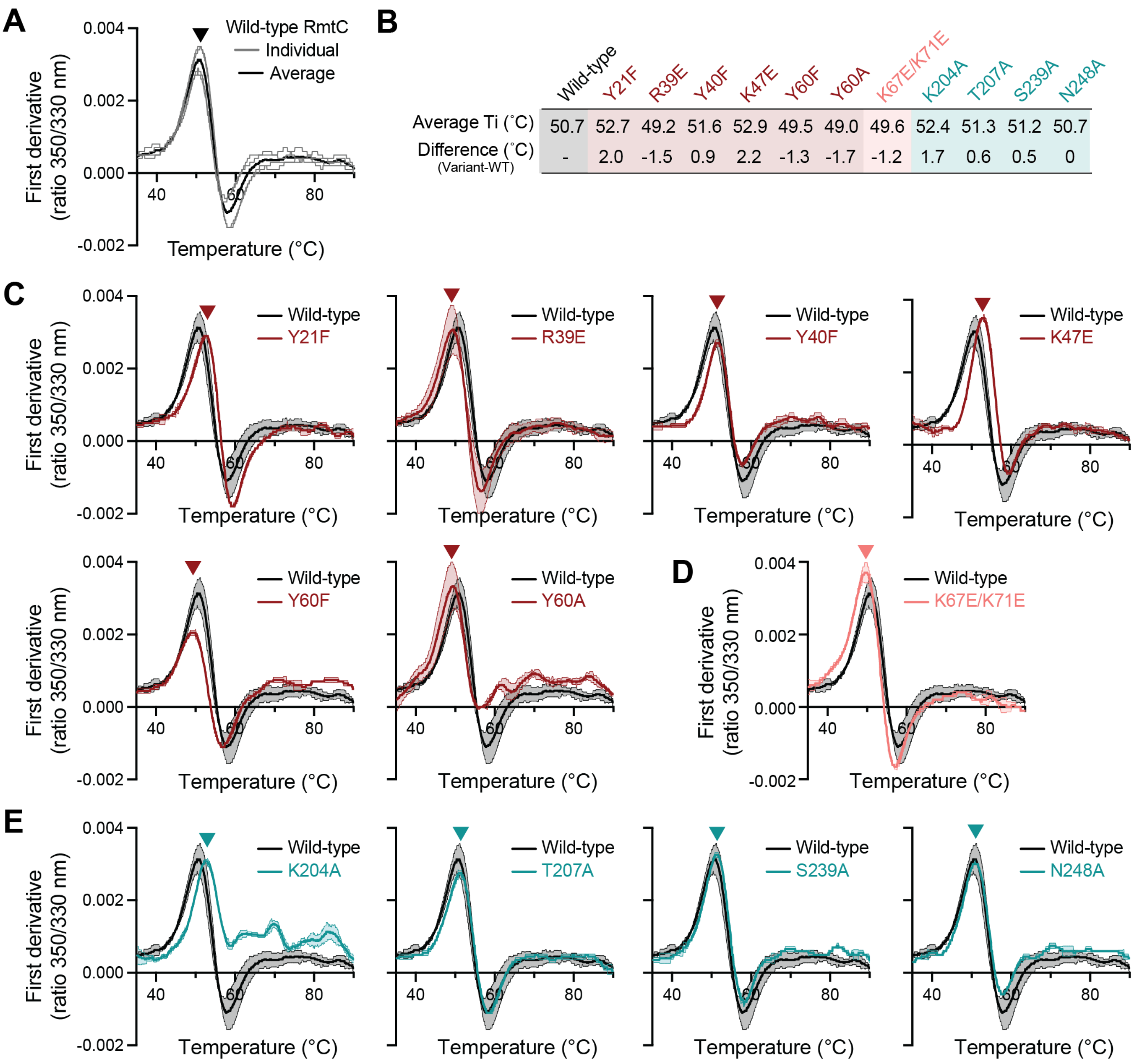


| | Wild-type | Y21F | R39E | Y40F | K47E | Y60F | Y60A | K67E/K71E | K204A | T207A | S239A | N248A |
|---|---|---|---|---|---|---|---|---|---|---|---|---|
| Average Ti (˚C) | 50.7 | 52.7 | 49.2 | 51.6 | 52.9 | 49.5 | 49.0 | 49.6 | 52.4 | 51.3 | 51.2 | 50.7 |
| Difference (˚C) (Variant-WT) | - | 2.0 | -1.5 | 0.9 | 2.2 | -1.3 | -1.7 | -1.2 | 1.7 | 0.6 | 0.5 | 0 |

**Figure S10**. **Quality control of purified wild-type and variant RmtC proteins by thermal denaturation. A**. Replicate measurements (gray lines) and average (black line) of wild-type (WT) RmtC unfolding monitored using intrinsic fluorescence at 330 and 350 nm (Tycho analysis), illustrating the reproducibility of the method between experiments and protein preparations. First derivative plots are shown for fluorescence ratio (350/330 nm) from which the inflection temperature ($T_i$) is derived (shown for the positive peak, marked with an arrowhead). **B.** Summary of Ti values for each protein and the calculated difference compared to the wild-type protein. Tycho analysis of variants generated in this study in the **C**. N1 subdomain (variant in dark red), **D**. N2 subdomain (light red, shown for the double substitution), and **E**. CTD (teal). The shaded regions on individual curves show the standard deviation between measurements.

**Supplementary Table 1: Conservation of select residues among m$^7$G1405 (16S rRNA) methyltransferases.**

| Residue | Residue identity (% occurrence)[a,b] | | |
|---|---|---|---|
| | All | Intrinsic | Acquired |
| K20 (N1) | K (69.1), R (25.0) | R (100) | K (83.3), N (11.1) |
| Y21 (N1) | Y (94.1) | Y (100) | Y (88.9) |
| R39 (N1) | R (47.8), K (37.3) | R (81.8) | R (55.6), K (44.4) |
| Y40 (N1) | G (26.9), R (19.4), H (17.9), Y (13.4) | G (100) | Y (44.4), H (33.3), F (16.7) |
| K43 (N1) | K (58.8), R (4.4) | NC | K (94.4) |
| K47 (N1) | K (79.4), R (5.9) | K (100) | K (61.1), E (33.3) |
| R50 (N1) | R (50.0), K (48.5) | K (100) | R (77.8), K (22.2) |
| H54 (N1) | H (98.5) | H (100) | H (100) |
| Y60 (N1) | Y (69.1), F (23.5), H (7.4) | Y (54.5), F (45.5) | Y (61.1), F (38.9) |
| K67 (N2) | NC | NC | K (66.7), R (16.7) |
| R68 (N2) | NC | NC | K (66.7), R (16.7) |
| K71 (N2) | NC | A (54.5), K (27.3) | R (27.8), K (11.1), A (38.9) |
| K72 (N2) | NC | R (54.5), G (36.4) | NC |
| S107 (N2) | S (98.5) | S (100) | S (94.4) |
| D160 (CTD) | D (98.5) | D (100) | D (100) |
| K204 (CTD) | K (100) | K (100) | K (100) |
| T207 (CTD) | P (76.5), H (17.6), T (5.9) | P (100) | P (88.9), T (11.1) |
| R211 (CTD) | Q (44.1), R (20.6), T (10.3) | T (63.6), R (18.2) | R (61.1), Q (16.7), K (5.6) |
| K236 (CTD) | R (38.2), K (23.5), H (10.3) | K (81.8), R (18.2) | R (61.1), K (38.9) |
| S239 (CTD) | G (66.2), S (19.1) | G (100) | G (72.2), S (27.8) |
| R241 (CTD) | R (77.3), K (18.2) | R (90.9), K (9.1) | R (83.3), K (16.7) |
| M245 (CTD) | M (76.5), L (11.8) | M (100) | M (88.9), L (11.1) |
| N248 (CTD) | N (32.4), T (14.7), R (16.2), H (11.8) | T (63.6), N (36.4) | N (44.4), F (11.1), H (33.3) |

[a]m$^7$G1405 (16S rRNA) methyltransferases from aminoglycoside-producing bacteria (intrinsic), pathogens (acquired) and all enzymes.

[b]NC indicates not conserved (less than 60% sequence identity and dissimilar properties with the enzyme group).
